# Whole Genome Bisulphite Sequencing Using the Illumina Hiseq X System

**DOI:** 10.1101/193193

**Authors:** M. Suzuki, W. Liao, F. Wos, A.D. Johnston, J. DeGrazia, J. Ishii, T. Bloom, M.C. Zody, S. Germer, J.M. Greally

## Abstract

The Illumina HiSeq X platform has helped to reduce the cost of whole genome sequencing substantially, but its application for bisulphite sequencing is not straightforward. We describe the optimization of a library preparation and sequencing approach that maximizes the yield and quality of sequencing, and the elimination of a previously unrecognized artefact affecting several percent of bisulphite sequencing reads.

---

While the comprehensive representation of the majority of the genome by whole genome bisulphite sequencing (WGBS) makes it the optimal assay for testing DNA methylation^1^, up to now its cost has made it too expensive for many projects. The release of the Illumina HiSeq X reduced the cost of whole genome sequencing (WGS) substantially, prompting us to develop a new protocol based on this instrument to reduce the cost of WGBS to a comparable extent.

The cost efficiency of the X system in part depends upon the libraries having large inserts that allow 150 bp paired end sequencing to work effectively without a high fraction of overlapping reads, a practical problem when using DNA treated with sodium bisulphite, which has a degradative effect. Illumina provides the TruSeq DNA Methylation Kit for WGBS, which uses post-bisulphite adaptor tagging (PBAT)^2^, and recommends using 75 bp paired end sequencing, suitable for the shorter fragments from PBAT libraries. Apart from the insert size issue, the use of patterned flow cells and different base calling software on the X system makes a transition from the use of earlier technologies potentially problematic, requiring the optimization of both library preparation and sequencing, as we describe below.

We developed a new transposase-based approach that we call BS (bisulphite)-tagging, illustrated in Figure 1. In its use of transposases, the assay resembles prior tagmentation-based bisulphite library preparation assays^3,4^, while the incorporation of 5mC for end repair is comparable with the T-WGBS approach used for very low input DNA amounts^5^, and creates an initial normal complexity sequence before reading into the (G+C)-depleted bisulphite-converted insert. Unlike T-WGBS, the extra expenses of a modified transposase and pre-methylated oligonucleotides are not required for BS-tagging.

**Figure 1.**
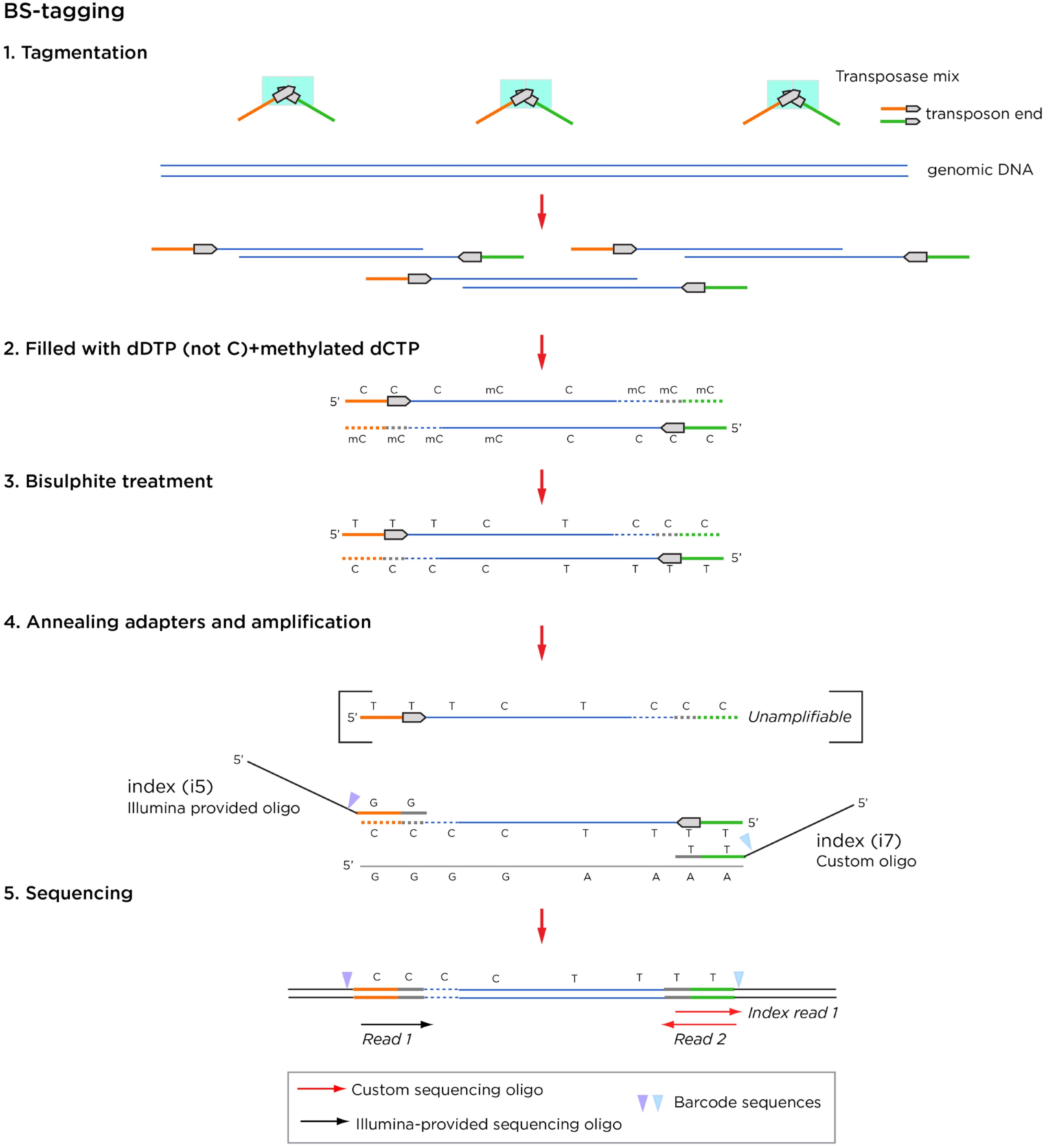
Overview of the BS-tagging assay. Standard transposases are used but with a 5mC fill-in. The bisulphite treatment changes the original transposon sequence and prevents it from being amplified, but the fill-in complement with 5mC remains amplifiable. One custom oligonucleotide is needed for this assay.

There are three types of duplicate sequences that can be generated using the X system. The X system-specific problem, because of the use of patterned flow cells, is when a library fragment occupying one well migrates or jumps into an adjacent well, referred to as a proximal duplicate. The second is the PCR duplicate, in which the same library fragment is amplified and is sequenced in different wells, while the third is the separate amplification of each of two complementary strands of DNA (complementary strand duplicate). Duplicates are usually identifiable by having the same start and end positions in the genome, allowing their removal by data filtering. As BS-tagging can only amplify one of the two complementary strands, these should never enter the library to begin with, reducing this source of non-productive sequencing. We show the duplicate rates for X-WGBS as part of **Supplementary Table 1**. Our stringent removal of smaller insert size library components in order to minimize overlapping read pairs (and hence effective coverage) reduces the amount of starting material, potentially contributing to the increased PCR duplicate rates compared with WGS, but without the additional penalty of complementary strand duplicates. Insert size optimization was achieved by adjusting transposase treatment and bead clean up conditions.

The sequencing was also customized. Early versions of the Illumina software^6^ (*e.g.* HiSeq Control Software v3.3.39, RTA v2.7.1) were not designed to handle unbalanced libraries and required a substantial spike-in of PhiX to generate data of reasonable quality. In that setting, we tested whether we could use an alternative source of spike-in DNA with a higher (G+C) content as a more effective balance for the (A+T)-rich bisulphite-converted DNA. We compared Illumina’s 44% (G+C) PhiX spike-in^7^ with a library from *Kineococcus radiotolerans*, which has 74% (G+C)^8^, and found the *K. radiotolerans* spike-in to perform better than PhiX when both were added at ∼17% proportions, and that even 5.3% *K. radiotolerans* was enough to restore base quality (Supplementary Fig. 1). Recent versions of HiSeq X software (RTA 2.7.7 and HCS 3.4.0.38) included a revised Q-table to facilitate sequencing of unbalanced libraries (including WGBS), allowing a 5% PhiX spike-in to generate high-quality sequencing data. However, since PhiX libraries are often slightly different in size than WGBS libraries, and the ExAmp process on the HiSeq X preferentially clusters smaller fragments, it is often difficult to achieve precise representation of PhiX spike-in in pooled libraries. A (G+C)-rich spike-in such as from *K. radiotolerans* in the setting of the new HiSeq X software is much more robust to such variation, with sequencing base quality maintained even at 2-3% spike-in levels. We developed a custom indexing primer that allows multiplexing of samples on the X system, including the use of a custom *K. radiotolerans* spike-in sample. As a control for bisulphite conversion, we used unmethylated lambda DNA as a second spike-in sample.

We show our analytical pipeline in Supplementary Fig. 2, and typical results in **Supplementary Table 1**. We compared the results of these X-WGBS data with DNA methylation data generated by other approaches in the same cell types. In Supplementary Fig. 3 we show the root-mean-square error (RMSE) values for X-WGBS, WGBS using older sequencing technology^9^, reduced representation bisulphite sequencing (RRBS), the SeqCap Epi capture approach that we developed^10^, and microarray-based assays (Infinium HumanMethylation450, Infinium MethylationEPIC, Illumina). The DNA methylation patterns of each cell type clustered separately, including the WGBS and X-WGBS results.

**Figure 2.**
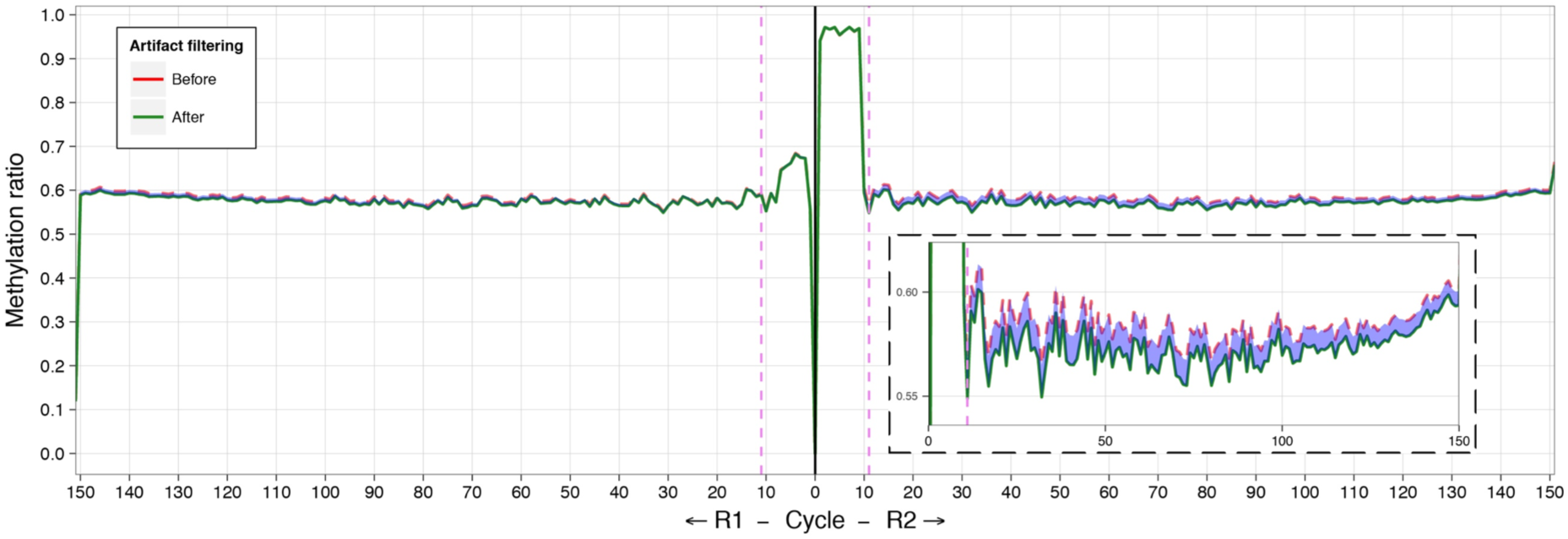
A subset of reads shows apparent cytosine methylation throughout their lengths, in CG and CH dinucleotide contexts. Our use of 5mC for end repair reveals the expected ∼10 bp increase in the C/T ratio (y axis) in read 1 (right). However, prompted by our finding of unconverted cytosines in lambda DNA, we tested whether any of the reads had a pattern of unconverted cytosines in all dinucleotide contexts (the expected CG and the uncommon CH contexts). We found a subset of reads in which no cytosines were converted. To categorize these, we developed the *filterFillIn2* software to identify four consecutive unconverted cytosines in a CHH context. The effect on DNA methylation values is shown in the C/T plots, the original values in red and following removal in green. The inset shows that ∼1-2% of the DNA methylation value is accounted for by this artefact.

A major rationale for the use of WGBS as opposed to survey assays is the ability to measure DNA methylation at distal *cis*-regulatory sites in diverse tissue types^1^. We tested relative performance by performing the Assay for Transposase-Accessible Chromatin with high throughput sequencing (ATAC-seq)^11^ as well as using data from the ENCODE project to define the *cis*-regulatory landscape in the GM12878 cell line. The loci of transposase-accessible chromatin coincide with loci of decreased DNA methylation and are flanked by enrichment for histone H3 lysine 4 tri- and monomethylation (H3K4me3, H3K4me1) and histone H3 lysine 27 acetylation (H3K27ac, Supplementary Fig. 4). The X-WGBS assay reports the large majority of loci of transposase-accessible chromatin in GM12878 cells, whereas RRBS is especially poorly representative of distal (non-promoter) regulatory loci (<10%) (Supplementary Fig. 5). The SeqCap Epi CpGiant design has approximately equivalent coverage of *cis*-regulatory sites as the Infinium HumanMethylation450 microarray, which reflects how both are designed to target the same loci. The Infinium MethylationEPIC microarray, designed to cover more sites than the HumanMethylation450 design, interrogates over half the ATAC-seq peaks overall and just under half of the distal *cis*-regulatory loci in GM12878 cells.

We noted the puzzling finding that lambda DNA appeared to have some unconverted cytosines indicating DNA methylation, which should not be occurring in this organism. When we explored this finding, we found the lack of conversion to occur in both CG and CH contexts throughout a minority of sequence reads. This indicates that our fill-in reaction using 5mC following transposase activity was extending beyond the typical several nucleotides in this subset of molecules. When we then explored the human DNA being simultaneously sequenced, we found that 1-2% of the reads showed a similar pattern (Fig. 2). This artefact is likely to have occurred in previous bisulphite sequencing reactions but remained unrecognized because of the use of unmodified cytosine in the fill-in reaction. We therefore developed an algorithm to identify 4 consecutive, unconverted CHH sites in a sequence read, allowing us to filter those for which we have a high suspicion of this technical artefact.

We show that the X-WGBS data reveal the many types of events typically sought in a study of mammalian DNA methylation. In **Supplementary Fig. 6** we show examples of low-methylated regions (LMRs)^12^, and differentially-methylated regions (DMRs)^13^. We also show that the XWGBS reads can be used to identify allele-specific DNA methylation^14^.

When we compare the reagent costs for the equivalent amount of WGBS on the X and HiSeq 2500 platforms, the relative cost reduction associated with X-WGBS is ∼4-fold compared to both HiSeq 2500 (2x125 bp) and 4000 (2x75 bp). This assumes 100 Gb raw data on the 2500 and 4000 at list prices (currently $32.60 and $30.24 per Gb respectively), and 120 Gb on the HiSeq X to account for the additional spike-in needed, as well as the elevated platform-specific duplicate rates. Library preparation costs for X-WGBS are similar to standard commercial kits. The similarity of the newer NovaSeq systems to the X technology in terms of patterned flow cells and chemistries indicates that migrating the assay to NovaSeqs may allow further cost reductions, assuming the RTA software is similarly tolerant of unbalanced libraries as recent versions of the HiSeq X software. Furthermore, as the BS-tagging library preparation approach should be highly amenable to automation, scaled use of the assay should realize still further savings. This decrease in the cost of the WGBS assay now allows it to be considered as a first line approach in population studies associating DNA methylation changes with phenotypes. We conclude that while adapting WGBS to the HiSeq X system created unique challenges, these can be overcome with BS-tagging, allowing more cost-effective mammalian WGBS than previously possible.

## Methods

Methods and associated references are available in the online version of the paper. The BS-tagging assay is described comprehensively in the **Protocol Exchange** resource (submitted) and the **Supplementary Data**.

## Acknowledgements

We are grateful to Einstein’s Center for Epigenomics, and the New York Genome Center (NYGC) personnel Stefan Pescatore and Harold Swerdlow, and the NYGC production sequencing team for their support.

## Author Contributions

Project design MS, SG, JMG. BS-tagging assay MS, FW. Experiments MS, FW, ADJ, JD. Analysis WL, TB, MCZ. Project management MS, JI, TB, SG, JMG.

## Supplementary Information

**FIGURE S1:**
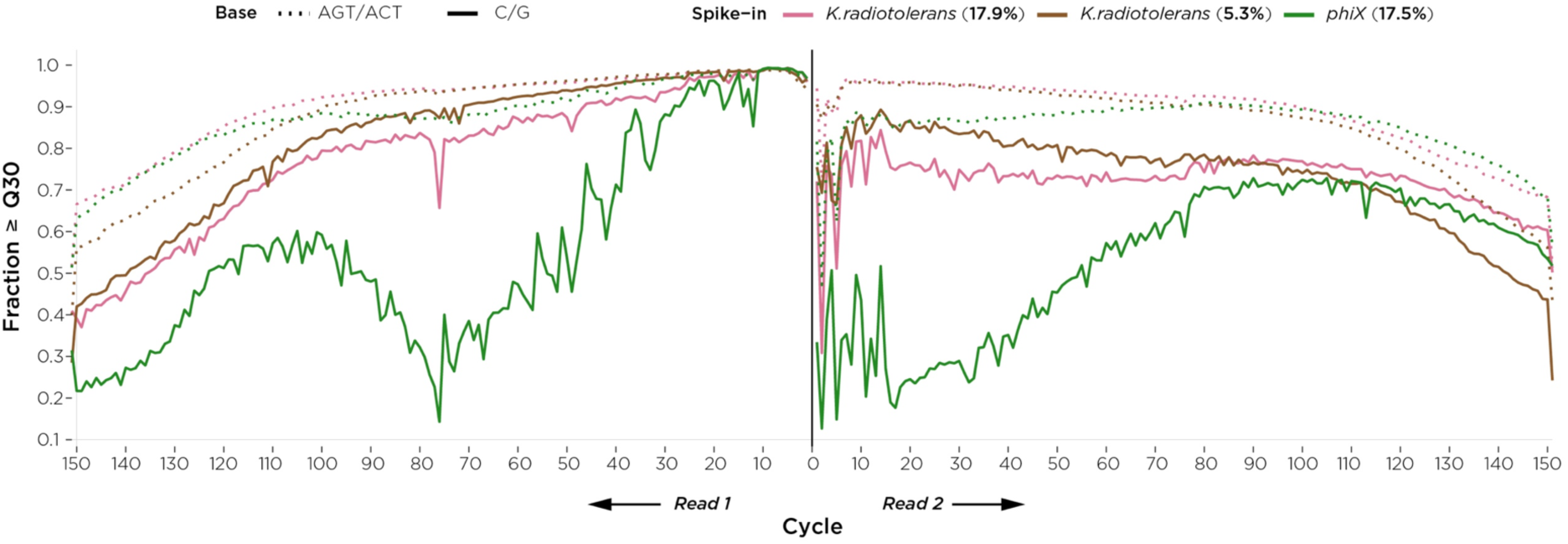
Improved quality of whole genome bisulphite sequencing (WGBS) with *Kineococcus radiotolerans* used as a spike-in. The plot shows the proportion of reads at or above a quality score of 30 for read 1 (left) and read 2 (right). The pink lines show the performance of *K. radiotolerans* (G+C=0.74) added at a proportion of 17.9% (light pink) or 5.3% (dark pink), and phiX (G+C=0.44, green). The results are plotted to show the quality for C/G (solid lines) separately from other nucleotides (dotted lines), as the performance of WGBS is especially problematic for C/G nucleotides, which is missed by plots that do not separate out these nucleotides. What is apparent is that the high (G+C) *K. radiotolerans* spike-in, even at 5.3% proportion, restores C/G quality to levels comparable with the other nucleotides.

**FIGURE S2:**
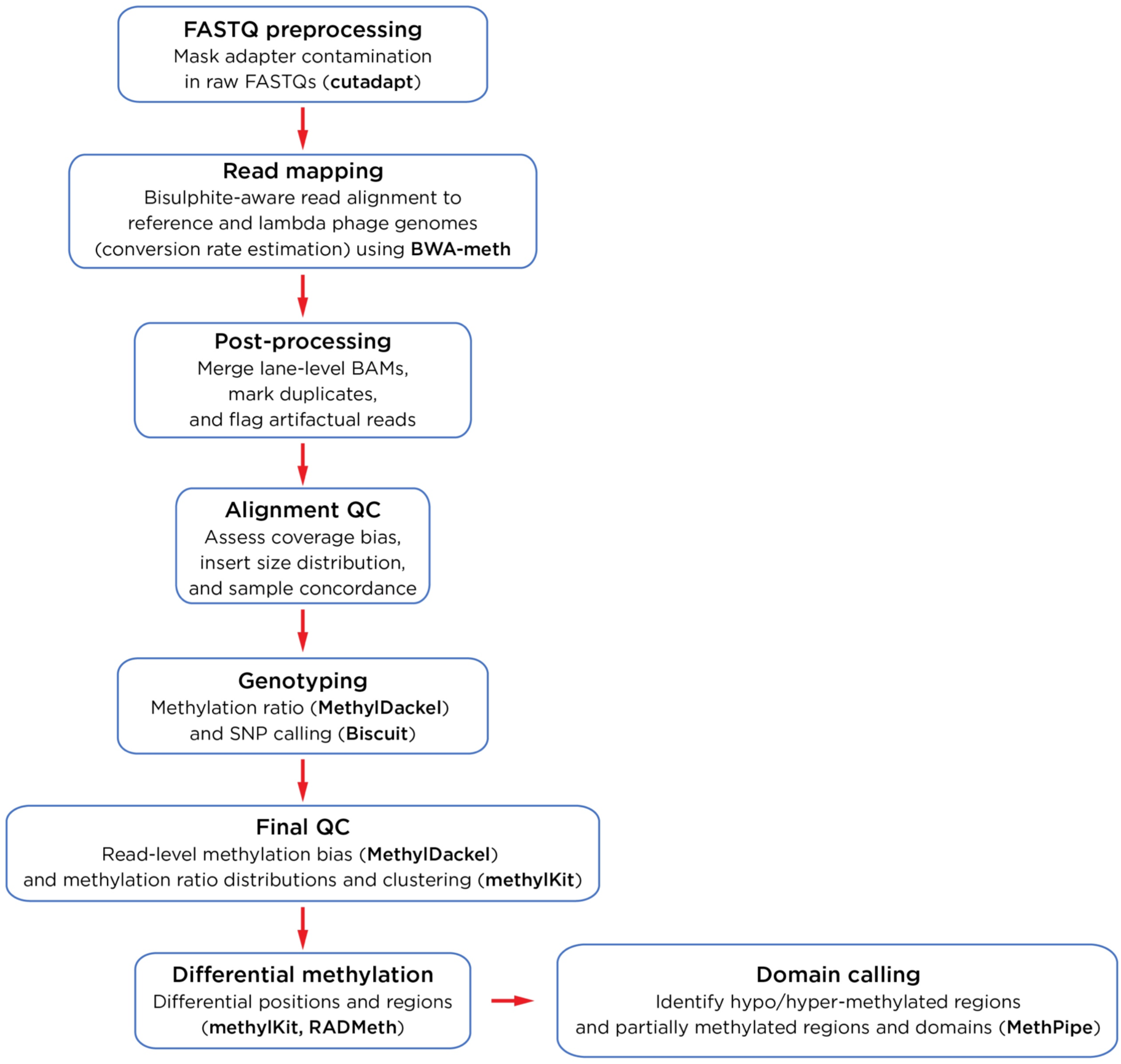
Overview of our analytical pipeline. The analytical pipeline we use is relatively standard for bisulphite sequencing analysis. The choices of individual analytical tools within the pipeline are highlighted.

**FIGURE S3:**
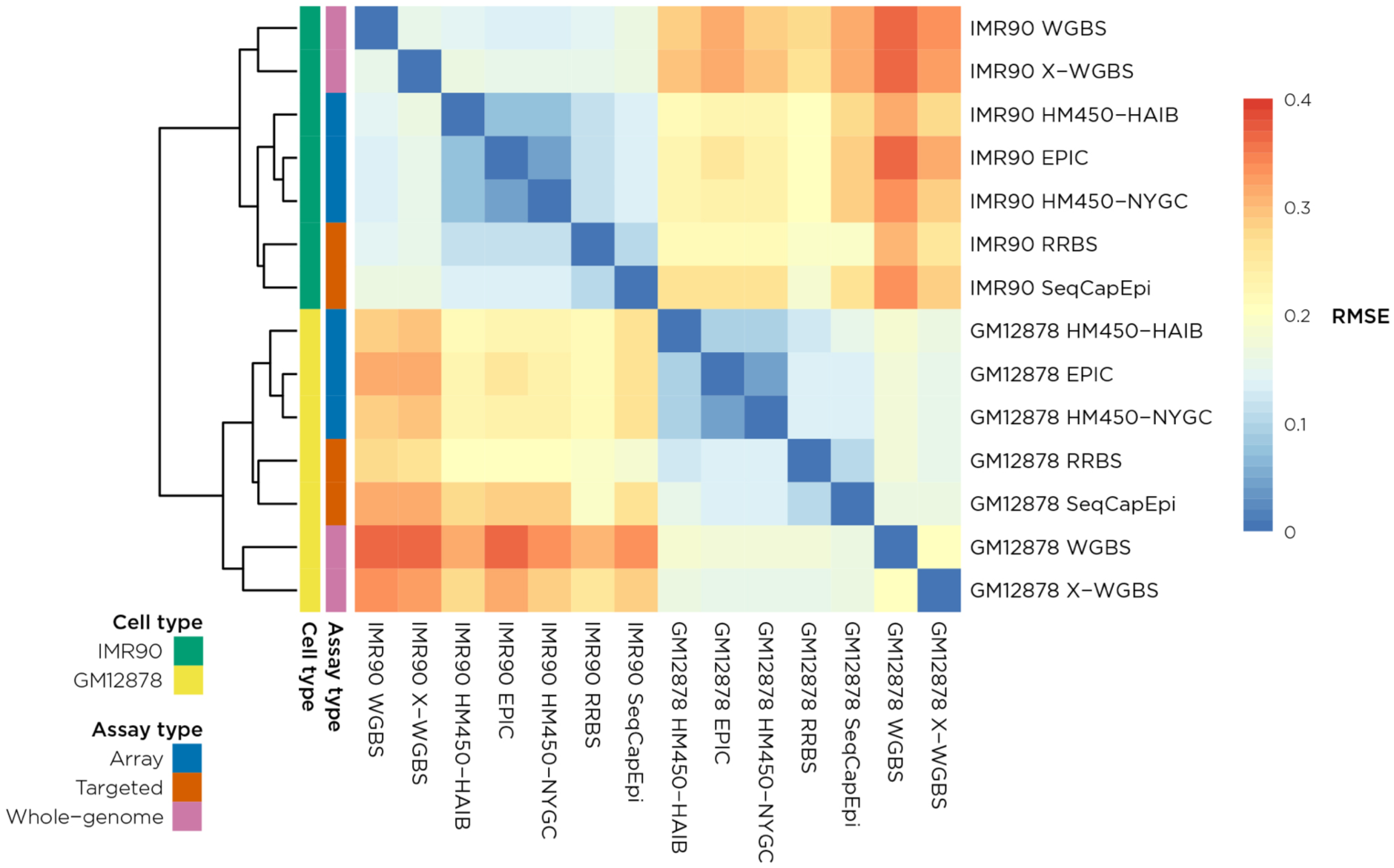
Concordance of DNA methylation measured by different assays in two cell types. We measure the concordance of DNA methylation values between samples using the root-mean-square error (RMSE), with lower error values (blue) indicating higher concordance between measurements. There is a clear difference between the values for the IMR90 and GM12878 cell types, with X-WGBS clustering with WGBS results using older technology.

**FIGURE S4:**
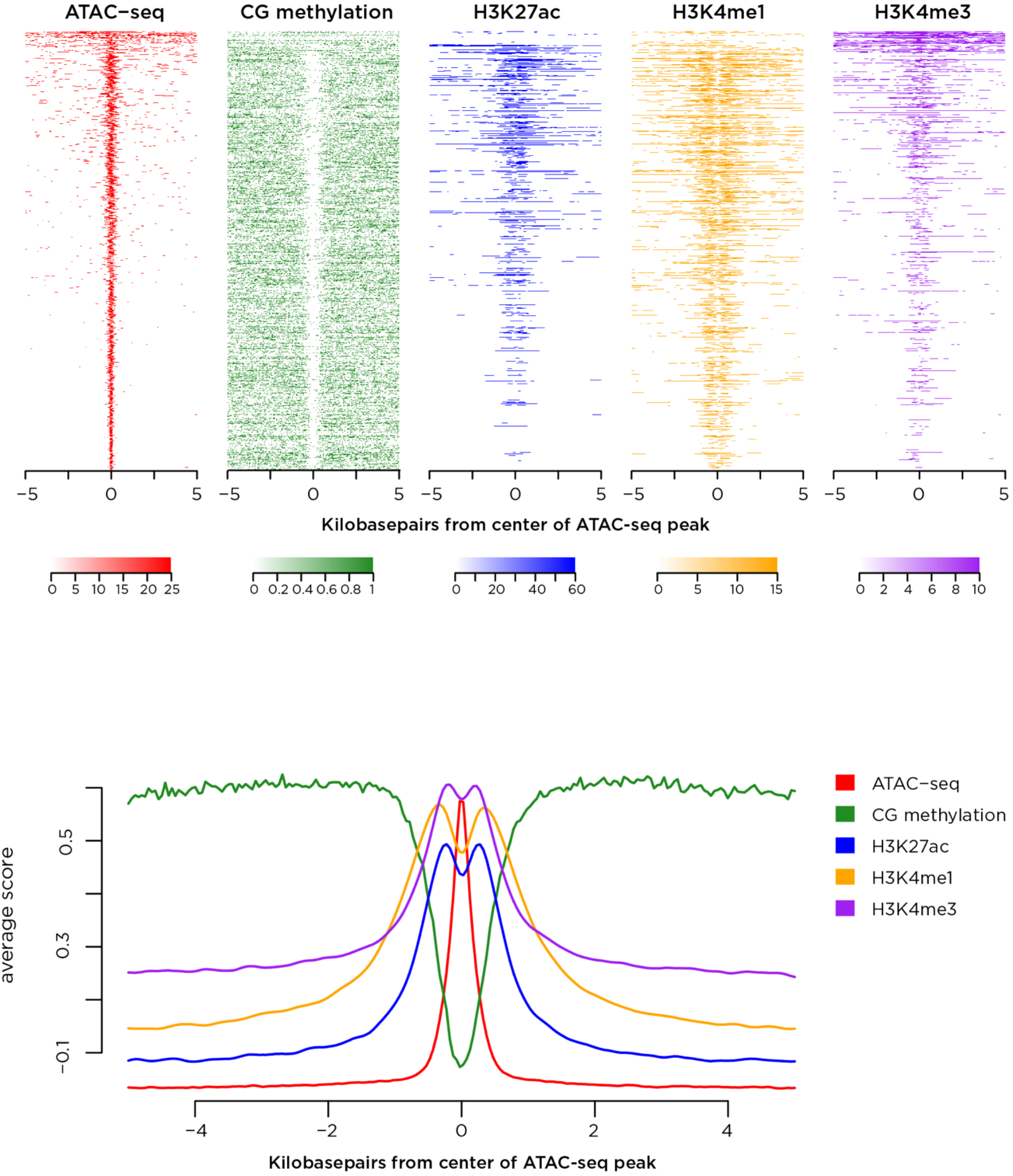
The loci of transposase-accessible chromatin correlate with hypomethylated DNA and chromatin modifications of active *cis*-regulatory elements. We used ATAC-seq to identify transposase-accessible chromatin, defining usually <1 kb regions (top left, red). The same loci show decreased DNA methylation (top, green), and local enrichment of post-translational modifications of histones, including H3K27c (blue), H3K4me1 (gold) and H3K4me3 (purple). These histone modifications are enriched one each side of the nucleosome-free regeion defined by ATAC-seq, better demonstrated in the lower plot.

**FIGURE S5:**
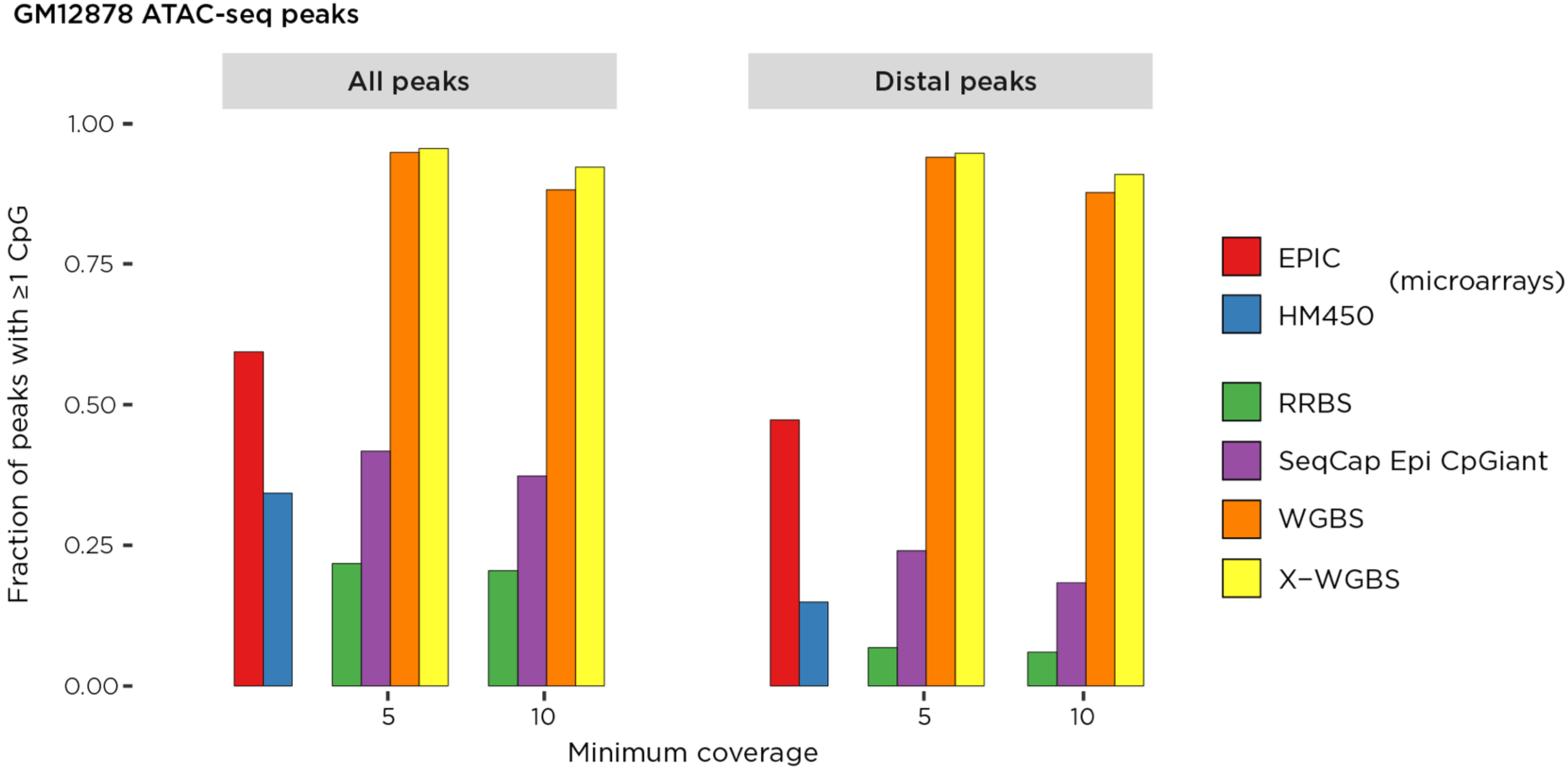
The relative representation of *cis*-regulatory elements genome-wide by different DNA methylation assays. We use the ATAC-seq peaks as a representation of the *cis*-regulatory elements in GM12878 cells. These were further subdivided into those distal (>10 kb upstream or >2 kb downstream) from RefSeq genes. The X-WGBS results interrogate at least one CG in the majority of these loci at 5 or 10X coverage, whereas RRBS represents only a small proportion of these loci, with the Infinium HumanMethylation450 microarray and the SeqCap Epi CpGiant system representing similar proportions (as expected for designs targeting the same genomic loci), with the expanded Infinium MethylationEPIC microarray representing a higher proportion of ATAC-seq peaks.

### Supplementary Methods

#### BS-tagging protocol

##### Step 1: Tagmentation of genomic DNA

This step generates fragmented DNA by tagmentation. The ratio of Tn5:DNA determines fragment size. Fluorometric measurement of input DNA should therefore be performed to quantify input DNA.

1. Set up the following tagmentation reaction in a deep well MIDI plate, and incubate at 37°C for 5 minutes in a Hybex oven:

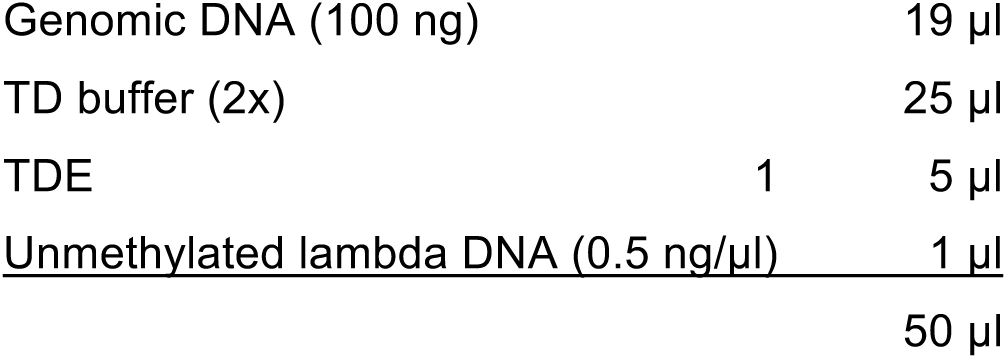 Gently pipette up and down 10 times to mix. Change tips between samples. Alternatively: mix on a thermomixer at 1,500 RPM for 30 seconds. Spin briefly prior to 37 ^o^C incubation. Change tips between samples. 2. Once the 5 minute incubation is complete, place the MIDI plate on ice and proceed immediately to **Step 2**.

##### Step 2: Silane bead reaction cleanup

1. To each well of the 96 well MIDI plate containing the tagmented product, add the following:

2. The total reaction volume will be 265 μl. Seal the plate and mix well by placing the MIDI plate on a thermomixer and shaking at 1,350 RPM for 2 minutes, then incubate the plate at room temperature for 5 minutes.
3. Place the plate on a magnetic stand for 3 minutes or until clear.
4. Carefully remove and discard the supernatant.
5. Remove the plate from the magnet and add 200 μl of 80% ethanol*
6. Seal the plate and mix well by placing the MIDI plate on a thermomixer and shaking at 1,350 RPM for 30 seconds.
7. Place the MIDI plate back onto the magnet for 2 minutes or until clear.
8. Carefully remove and discard the supernatant when clear and proceed to repeat steps 5-7 for a total of two ethanol washes.
9. After the second wash, place the plate on magnet for 2 minutes or until clear.
10. Remove and discard the supernatant then briefly spin down to collect any residual ethanol at the bottom of the well.
11. Place the plate back on the magnet and use pipette to remove any remaining ethanol.
12. Dry the beads at room temperature for 4-6 minutes.
13. Once dry, remove the MIDI plate from the magnetic stand and elute in 21 μl elution buffer.
14. Seal the plate and mix by shaking at 1,850 RPM for 4 minutes.
15. Briefly spin the plate and place it back onto a magnetic rack for 3-5 minutes or until clear.
16. Carefully transfer 19 μl of the eluted material into a clean PCR plate.
17. Proceed immediately to **Step 3**.

|  |  |
| --- | --- |
| i. Dynabeads® MyOne™ Silane | 15 µl |
| ii. <u>Qiagen Buffer PB</u> | <u>200 µl</u> |
|  | 215 µl |

* Use freshly prepared 80% ethanol for the washing step

##### Step 3: End -filling with 5-methyl-dCTP mix

1. Set up end filling reaction in a clean PCR plate and incubate at 37 ^o^C for 30 minutes in a thermal cycler:

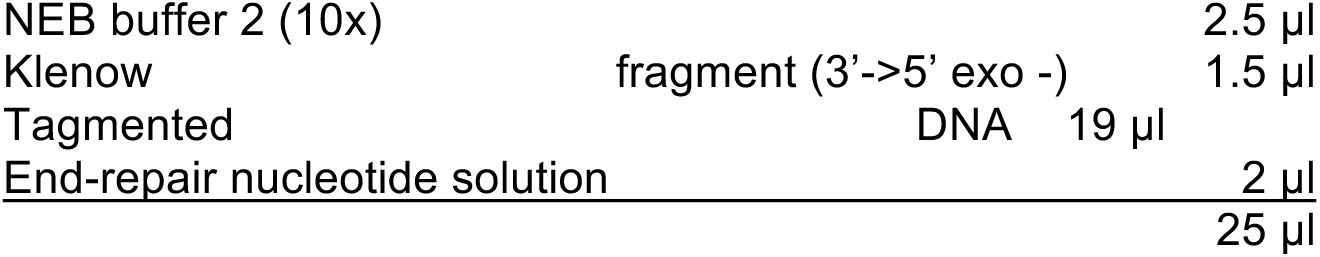
2. Clean up the product with 50 μl (2:1 ratio) of AMpureXP beads according to the manufacturer’s instructions. The elution volume is 21 μl in elution buffer.

##### Step 4: Bisulphite treatment

1. Perform the bisulphite treatment reaction according to the manufacturer’s instruction. **CRITICAL STEP** | ensure that the conversion mix is performed for at least 10 minutes while protected from light. There should be no visible solids in the conversion reagent. If solids are present, a fresh conversion mix should be made to avoid conversion issues.
2. The final elution volume is 18 μl

##### Step 5: Adapter attachment and PCR amplification

1. Set up the following reaction in a PCR plate:

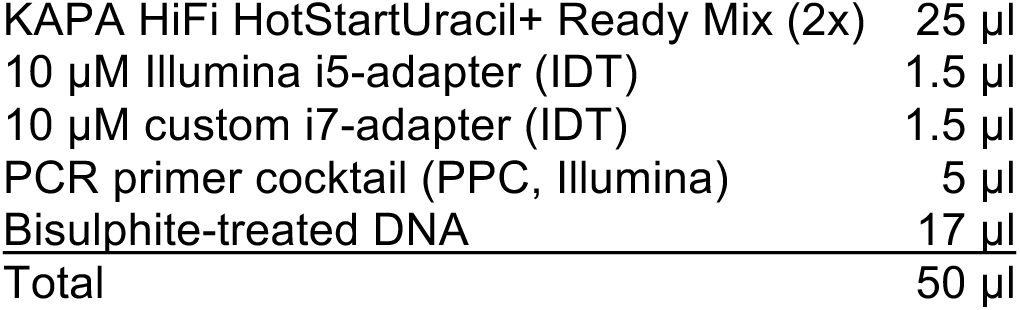
2. Place the sample plate in a thermal cycler and perform the following steps:

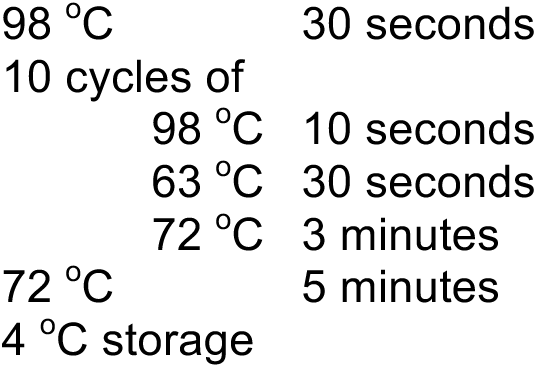

##### Step 6: Size selection of library

1. Set up the purification reaction in a deep well, MIDI plate and purify the products according to the manufacturer’s instructions:

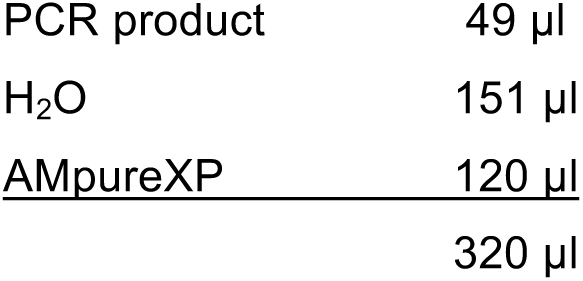 * Use freshly prepared 80% ethanol for the washing step. **CRITICAL STEP** | The use of a 0.6X AmpureXP cleanup will exclude fragments <400 bp
2. Elute in 15 μl of elution buffer.

##### Step 7: Library quality check

Before running the libraries on a massively parallel sequencer, check the length distribution of library using the Bioanalyzer and quantify the product with Qubit Fluorometric Quantitation.

##### Step 8: Massively parallel sequencing on the Illumina HiSeq X system

Custom sequencing primer oligos should be used for read 2 sequencing and i7 index sequence reading. The Illumina-provided sequencing oligos are used for read 1. If using recent versions of HiSeq X software (RTA 2.7.7 and HCS 3.4.0.38), a spike-in of 5% PhiX DNA should be enough to generate high-quality data, but an even lower proportion of *K. radiotolerans* should generate data of equivalent quality.

##### Stage 9: Analyzing BS-tagging data

Adapter sequence are first N-masked from raw FASTQ files using *cutadapt* v1.9.1^1^. Because the library protocol produces R1 reads with G>A transitions and R2 reads with C>T transitions after bisulphite conversion, short-read alignment is performed with *bwameth*^2^ with R2 reads mapped as a typical C>T converted R1 read and R1 reads mapped as a typical G>A converted read. Additionally, we modified *bwa-meth*’s default minimum longest match length for a read (0.44*read-length) to greater than 30 bp (0.2*read-length) before marking the alignment in sequence alignment/map (SAM) format^3^ with the 0x200bitwise flag indicating not pass filter, failed platform/vendor quality control. The resulting alignments are marked for duplicates using *Picard* v2.4.1. To calculate strand-specific duplicates, alignments are separated based on their orientation to the reference, then *MarkDuplicates* is run on the two alignment sets separately. Our use of methylated cytosines in end-repair following tagmentation revealed a rare artifact (∼1-2% of reads) where incorporation of the methylated cytosines extended deeper into the duplex fragment than the typical ∼9 bases expected from the transposition footprint. Because this obscures the original methylation state, we designed a custom Perl script, *filterFillIn2,* which marks reads with 4 consecutive methylated CHH’s outside of the first 9 bases and excludes those reads from downstream analysis by appending the 0x200 bitwise flag, marking them as qc failed.

#### DNA methylation microarray analysis

Illumina Infinium HumanMethylation450 (HM450) and MethylationEPIC (EPIC) arrays performed at NYGC were prepared and processed according to manufacturer specifications. Raw IDAT files were processed to Beta values using the *minfi* R package^4^ using *illumina* background correction and subset within array normalization (*SWAN*)^5^ to correct for probe type bias.

#### ATAC-seq library preparation

The assay for transposase accessible chromatin (ATAC)-seq libraries were prepared similarly to Buenrostro *et al.*^6^. We used 50,000 cells from two biological replicates of GM12878, harvested during the exponential growth phase. The cells were spun at 500 g for 5 minutes at 4°C and then washed using 50 μL of cold 1X PBS. Samples were then centrifuged at 500 g for 5 minutes. Cells were lysed in cold lysis buffer (10 mM Tris-HCL, pH 7.4, 10 mM NaCl, 3 mM MgCl_2_ and 0.1% IGEPAL CA-630) and immediately spun at 500 g for 5 minutes at 4°C. The pellet was then resuspended in the transposase reaction mix (25 μL 2x TD buffer, 2.5 μL transposase and 22.5 μL nuclease-free water; Illumina Nextera). Following a 30-minute incubation at 37°C, the samples were purified using the Zymo DNA Clean and Concentrator purification kit. Following the purification, the libraries were amplified with the following PCR conditions:

> 72°C for 5 minutes;
>
> 98°C for 30 seconds;
>
> A total of 10 cycles of 98°C for 10 seconds; 63°C for 30 seconds; 1 minute for 72°C.

Subsequently, the libraries were purified using Agencourt AMpure XP beads; large fragments were filtered by using 0.6x (of PCR mixture volume) magnetic bead volume and taking the supernatant. Primer-dimer and short fragments were removed by collecting bead-associated DNA in a 1:1 (bead solution volume:mixture volume) mix. The two replicates were run on different Illumina HiSeq 2500 flowcells to obtain 100 bp paired-end reads, resulting in a mean of 57 million paired-end reads per sample.

#### ATAC-seq peak calling

Sequenced reads were aligned to the *Homo sapiens* (hg19) reference genome using the Burrows-Wheeler Aligner (*BWA-MEM* version 0.7.13)^3^. Reads were filtered using *picard-tools* v1.92 and *samtools* v0.1.19^7^ and shifted using *bedtools2* v2.24.0^8^ before peaks were called for each replicate using *MACS2* v.2.1.0^9^. Irreproducible discovery rates (IDRs) were found for the overlapping peaks using the method previously described^10^. Peaks with an IDR of less than 0.05 were filtered by consensus blacklisted regions and mitochondria homologous regions. Finally, the remaining peaks were retained for analysis.

#### Access and analysis of public data

Previously published whole genome bisulphite and SeqCap Epi data were downloaded from GEO (GSE16256) and DDBJ (SRP049215), respectively, and passed through the same analytical pipeline as our BS-tagging data, except without flipping R1 and R2 during alignment or excluding fill-in artefacts. For SeqCap Epi data, duplicates were also retained for downstream analysis. For the public RRBS and HM450 data, pre-processed methylation ratios were obtained from the UCSC Table Browser. Histone modification ChIP-seq datasets (GSE16256) were downloaded from the ENCODE/Roadmap data matrix (https://www.encodeproject.org/). Replicates were consolidated into a single high coverage dataset. FASTQs were adapter trimmed using *Trim Galore*!, then aligned to the hg19 reference genome using *BWA-mem*^3^. Alignments were further processed using the Genome Analysis Toolkit pipeline, performing insertion/deletion realignment and base quality score recalibration^11^. Duplicates were marked using Picard v2.4.1. Read fragment estimation was performed with *phantompeakqualtools* and used to extend alignments to estimated fragment size then piled-up, ignoring non-unique and duplicate alignments, to determine ChIP signal. Signals were normalized by library size (number or reads aligned to autosomal chromosomes) then scaled to the equivalent of 1X coverage using *deepTools*^12^.

#### Integrated data analysis

Comparison of DNA methylation ratios was performed at CpG dinucleotides covered at ^3^5X in the sequencing data. Heatmaps of RMSE values were generated and clustered by complete-linkage based on Euclidean distance (**Figure S3**). To assess various assay signals across putative regulatory regions defined by the ATAC-seq data (**Figure S5**), we focused only on distal accessibility peaks, defined as those ATAC peaks falling outside 10 kb upstream or 2 kb downstream of a gene (UCSC Known Genes canonical set). The ATAC-seq transposition signal was calculated based on read alignments centered at the putative transposase insertion point position using only R1 of the read pairs to prevent double counting of insertion instances. For signal heatmaps (**Figure S4**), we further restricted the analysis to ATAC peaks with an IDR score £0.016 to reduce the set to the top candidates that were of a more manageable size for plotting. Mean assay signal was obtained at 50 bp adjacent bins tiled across a region ±5 kb around summits of these selected ATAC-seq peaks. The WashU EpiGenome Browser (http://epigenomegateway.wustl.edu/browser/) was used to visualize user-provided and public datasets including the 15-state ChromHMM classifications^13^. Candidates for allele-specific methylation were selected based on previously implicated transversions^14^ and manually inspected for allele-specific methylation.

#### Computational requirements

The core analysis steps of pre-processing raw FASTQ files, bisulphite-aware short-read alignment, and DNA methylation genotyping were performed on the New York Genome Center (NYGC) high-performance compute cluster requiring roughly ∼300 core hours for a 30X bisulphite genome on Intel Xeon 2.60 GHz CPUs with >48 GB of available memory. Additional analysis time was required to perform various quality control steps, including but not limited to bisulphite conversion rate estimation, coverage uniformity assessment, and mean coverage estimation (both genome-wide and across *cis*-regulatory loci). Downstream analysis such as differential methylation detection, and hypo-/hypermethylated domain prediction required further computational time.

#### Data availability

All data generated are available through the Gene Expression Omnibus (GEO).

https://www.ncbi.nlm.nih.gov/geo/query/acc.cgi?acc=GSE103505

#### Code sources

*filterFillIn2* https://github.com/will-NYGC/bstag

*CutAdapt* https://cutadapt.readthedocs.io/en/stable/

*bwa-meth* https://github.com/brentp/bwa-meth

*Picard* http://broadinstitute.github.io/picard/

*Trim Galore*! https://www.bioinformatics.babraham.ac.uk/projects/trim_galore/

*Phantompeakqualtools* https://github.com/kundajelab/phantompeakqualtools

*deepTools* https://deeptools.readthedocs.io/en/latest/

Analytical pipeline modules (**Figure S2**):

*MethylDackel* https://github.com/dpryan79/MethylDackel

*MethPipe*^15^ https://github.com/smithlabcode/methpipe

*RADmeth*^16^ https://github.com/smithlabcode/radmeth

*methylKit*^17^ http://bioconductor.org/packages/release/bioc/html/methylKit.html

*Biscuit* https://github.com/zwdzwd/biscuit

